# Prevalence of Hepatitis B Virus (HBV) Basal Core Promoter/Precore region molecular variants among HIV/HBV co-infected and HBV mono-infected patients in Ile-Ife, Nigeria

**DOI:** 10.1101/2020.07.16.206151

**Authors:** Oluwadamilola Gideon Osasona, Lydia Boudarene, Opeoluwa Adewale-Fasoro, Uwem George, Judith Oguzie, Olumuyiwa Elijah Ariyo, Testimony J. Olumade, Awe Adeniyi, Moses Olubusuyi Adewumi, Adekunle Johnson Adeniji

**Author notes:** Corresponding author (O.G.O). Other authors (O.A.F), (U.G), (J.O), (T.J.O.), (O.E.A.), (L.B.), (A.A.), (M.O.A.), (A.J.A).

## Abstract

**Introduction:** Evolution of phenotypic diversity among viruses occurs as an escape mechanism against host immune pressure or drug selective pressure. Among HIV/HBV co-infected individuals, various HBV basal core promoter (BCP)/precore (PC) region molecular mutants had been reported with associated phenotypic defect in HBeAg production. The emergence of HBeAg negative variants of HBV in HIV co-infected individuals have profound implication on the diagnosis, management and prognosis of this subset of individuals. This includes delayed clearance of HBV, early development of adverse hepatic events such as liver cirrhosis and hepatocellular carcinoma. Currently, little is known about HBV BCP/PC region genomic heterogeneity in HIV/HBV co-infected patients in Nigeria. Therefore, this study was focussed on investigating evidence of precore/core region genomic variability among HIV/HBV co-infected patients in Nigeria.

**Materials and methods:** A total of 40 patients (20 HIV/HBV co-infected and 20 HBV mono-infected samples) were enrolled into the study and subsequently tested for HBsAg, HBeAg and HBeAb using specific Enzyme-Linked Immunosorbent Assay (ELISA). The BCP/PC genome regions (nucleotides 1653-1959) were amplified using a nested PCR assay and then subjected to BCP/PC mutational analysis in genome sites affecting HBeAg expression especially at the BCP transcriptional and PC Translational stop codon sites.

**Results:** Overall, 5(83.3%) of the six exploitable sequences after analysis showed various BCP/PC mutations. Only 1(16.6%) sequence from an HIV/HBV co-infected patient had the BCP transcriptional (double mutation; A1762T/G1764A) mutant. Analysis of the PC translational stop codon showed 4 (66.6%) having the G1896A mutants while 33.3% (2) had G1899A mutants.

**Conclusion:** This study has broadened the available evidence of BCP/PC region molecular mutants among HIV/HBV co-infected patients in Nigeria and assessed the difference of mutation prevalence in comparison with HBV mono-infected cohort. We therefore recommend that HIV/HBV co-infected patients be routinely screened for hepatitis B virus precore region mutants to improve their patient outcome.

## Introduction

Hepatitis B virus (HBV) infects about 2 billion individuals globally^1^ while about 350 million individuals have chronic HBV infection, and this accounted for mortality of 887,000 from hepatocellular carcinoma and liver cirrhosis in 2015 [1]. The bulk of these are from African and Asian continents [1]. About 38 million people globally are infected with HIV, and two-thirds of these are found in sub-Saharan Africa [2]. HBV and HIV have similar mode of transmission [2]. The global prevalence of HBsAg among HIV/HBV co-infected patients is 7.4%^2^. In HIV/ HBV co-infection, HBV enhances the progression of HIV, with worsening of immunologic and clinical parameters while conversely, HIV promotes the development of adverse hepatic events like cirrhosis and hepatocellular carcinoma [2].

The serological markers of HBV infection include Hepatitis B virus surface antigen (HBsAg), Hepatitis B virus surface antibody (HBsAb), Hepatitis B virus e antigen (HBeAg), Hepatitis B virus e antibody (HBeAb), Hepatitis B virus core IgG antibody (HBcIgG) and Hepatitis B virus core IgM antibody (HBcIgM). The HBeAg is a marker of viral replication and infectivity [3]. The development and eventual detection of HBeAg marker is often associated with active viral replication which is crucial in determining liver damage especially in chronic hepatitis B (CHB) viral infection. However, the disappearance of HBeAg and appearance of HBeAb is associated with reduced viral proliferation [3]. There have been documented evidence of HBeAg negative CHB infection which is often characterized by active liver diseases with persistently increased HBV DNA levels and active biochemical activity [4]. Studies have shown that most mutations that give rise to HBeAg negative variants arise from mutation in the precore region due to frameshift mutations or premature stop codon resulting in inability to produce the e antigen [5]. Also, some basal core promoter mutations that inhibit the production of precore mRNA at the level of transcription may also produce the HBeAg negative variants [5]. The most common mutation that accounts for the HBeAg negative variants is the G1896A – a point mutation involving change from G to A at nucleotide position 1896. This changes the 28^th^ codon of precore region from tryptophan (UGG) to a translational stop codon (UAG) with resultant premature termination of HBeAg translation process and inability to produce the viral HBeAg. The low levels of HBeAg in this category of patients may be characterized by alteration in HBV replication and liver diseases progression [6].

Little is known about HBV precore region genomic heterogeneity in HIV/HBV co-infected patients in Nigeria. In this study, we investigated the available evidence of the HBV precore/core gene variability amongst HIV/HBV co-infected patients and assessed the difference of mutation prevalence with a selected cohort of HBV mono-infected patients in Nigeria.

## Materials and methods

### Study Location and sample collection

This work is a cross-sectional hospital-based study carried out in the southwestern part of Nigeria, at the Virology clinic of the Obafemi Awolowo University Teaching Hospital, Ile-Ife. Ethical approval was granted by Obafemi Awolowo University Teaching Hospital, Ile-Ife and London School of Hygiene and Tropical Medicine Review Boards respectively. The study design conformed to the 2013 declaration of Helsinki. Socio-demographic data and clinical information were collected from each participant at the time of sample collection using self-administered structured questionnaire. Twenty HIV/HBV co-infected patients reporting at the virology clinic were recruited through purposive sampling technique involving screening of all HIV positive adult patients (250) for HbsAg. Twenty patients(8%) were found positive for HBsAg among the HIV positive patients, we also recruited same number (20) of known HBV mono-infected adult patients from the clinic who tested positive for HBsAg to have a basis for comparison of mutation rates in these two subgroups of patients. Study was conducted between May to August, 2019.

Five (5) millilitres of blood were collected from each participant by phlebotomist into labelled EDTA bottle using sterile needle and syringe after sterilizing the venepuncture site with 70 % ethanol soaked cotton wool. Blood samples were separated immediately by centrifugation at 1,500g for 15 min, and two plasma aliquots were aspirated into labelled sterile cryovials. The labelled cryovials containing plasma samples were transported on ice to the Department of Virology, College of Medicine, University College Hospital, Ibadan where they were preserved at -80°C until analysis.

### Laboratory Analysis

#### Detection of HBsAg, HBeAg and Anti-HBe Antibody Assay

All the 250 HIV positive plasma samples from patients enrolled into the study were initially screened for HBsAg using one step HBsAg strip (ACON Laboratories incorporated, USA) of which twenty(20) (8%) HIV positive patients tested positive for HBsAg. Another 20 HBV mono-infected patients (previously tested HBV positive at the centre) enrolled into the study were equally screened using one step HBsAg strip. All 40 plasma samples positive for HBsAg (including 20 plasma samples of HIV/HBV co-infected patients and 20 plasma samples of HBV mono infected patients) were subsequently subjected to HBsAg, HBeAg and HbeAb specific Enzyme-Linked Immunosorbent Assay (ELISA) (MELSIN Diagnostic kits China). Both the one-step HBsAg strip and ELISA assay were performed according to the manufacturer’s instructions. For the ELISA assay, optical density (OD) was read using the Emax endpoint ELISA microplate reader (Molecular Devices, California, USA) and the interpretation was made in line with the manufacturer’s instructions.

#### Nucleic acid extraction and HBV partial S-gene amplification

Total nucleic acid was isolated from all Hepatitis B surface antigen (HBsAg) ELISA positive plasma samples (n = 40) using Jena Bioscience viral RNA+DNA Preparation Kit. The extraction was done following the manufacturer’s instruction. The HBsAg specific polymerase chain reaction was carried out using a nested PCR (nPCR) targeting the partial S-gene region as previously described [7.8]. The first-round primers used were HBV_S1F (CTAGGACCCCTGCTCGTGTT),andHBV_S1R (CGAACCACTGAACAAATGGCACT), while the second-round primers were HBV_SNF (5-GTTGACAAGAATCCTCACAATACC-3) and HBV_SNR (5-GAGGCCCACTC CCATA-3). Both the first and second-round PCR reaction conditions were similar except that for first-round PCR, DNA extract from the sample was used as a template while first-round PCR product was used as a template for second-round PCR. PCR amplification was performed with a total volume of 50μL reaction containing 10μL of Red load Taq (Jena Bioscience, Jena, Germany), two microliters of each primer (made in 25μM concentrations), 4μL of DNA and 32μL of RNase free water. Thermal cycling was done using Veriti Thermal cycler (Applied Biosystems, California, USA.) as follows; 94°C for 3 minutes, followed by 45 cycles (denaturation; 94°C for 30 seconds, annealing; 55°C for 60 seconds and elongation; 70°C for 40 seconds with a ramp of 40% from 55°C to 70°C). After that, the reaction was further elongated at 72°C for 7 minutes and held at 4°C till terminated. Finally, PCR products were resolved on 2% agarose gel stained with ethidium bromide and viewed using a UV transilluminator.

#### HBV basal core promoter and precore (BCP/PC) region amplification

The BCP/PC genome regions (nucleotide (nt) 1653-1959) were amplified using a nested PCR assay and reaction mixture previously reported [9]. Briefly, 5 μL of the extracted DNA was mixed with 45μL of a PCR reaction mixture containing 400nM of the outer forward primers (5’-GCATGGAGACCACCGTGAC-3’)andreverseprimers(5’-GGAAAGAAGTCCGAGGGCAA-3’ in the first round PCR reaction. Nested PCR was done using forward primers (5’–CATAAGAGGACTCTTGGACT-3’ and reverse primers (5’-GGCAAAAAACAGAGTAACTC-3’) in the second round PCR reaction. The polymerase chain reaction thermal cycling condition was done as follows: 94°C for 30sec, followed by 30cycles (denaturation : 94°C for 30 sec, annealing: 55°C for 60 sec, elongation : 72°C for 40seconds), final extension: 72°C for 2 minutes. Reaction mixture concentration and conditions for PCR were similar for both the first and second-round PCR except that the cycles were increased to 35 cycles in the second-round amplification and first-round PCR products were used as template for second-round PCR. All PCR products were resolved on 1.5% agarose gel stained with ethidium bromide and viewed using a UV transilluminator.

#### Amplicon Sequencing

All PCR products that were positive for both HBV partial S-genes and BCP/PC specific polymerase chain reaction by having the required amplicon sizes(226 and 306 bp respectively)after viewing using a UV transilluminator were shipped to Macrogen Europe, Netherland for PCR product purification and BigDye chemistry sequencing. Second round primers for each assay were used for sequencing.

### Phylogenetic analysis and inference of Serotypes

The forward and reverse sequencing results per isolate for both S-gene and BCP/PC sequences were merged into contigs, and thereafter subjected to a BLASTn search on the NCBI BLAST webpage. GenBank sequences with high matching scores were retrieved and retained for downstream multiple sequence alignment for both genotyping and BCP/PC mutational analysis.

Phylogenetic analysis in this study was performed as previously described [10], and HBV serotypes were predicted based on analysis of amino acid residues at positions 122, 127, 134 and 160 specifying HBsAg determinants in the S region. Nucleotide sequences of S genes were retrieved from the HBV database (http://hbvdb.ibcp.fr/HBVdb/), and alignment done using the CLUSTAL W program in MEGA X software with default settings [11]. Phylogenetic tree was constructed in MEGA X software with the Kimura-2 parameter model [12,13] and 1,000 bootstrap replicates was used to construct a neighbour-joining tree.

### BCP/PC gene variability analysis

For BCP/PC mutational analysis, each sequence from this study were compared against known wild-type GenBank Genotype E reference sequences: X75657 and previously reported BCP/PC nucleotide sequences in Africa [9], using the CLUSTAL W program in MEGA X software with default settings [11]. Focus was given to major BCP/PC mutational genome sites affecting HBeAg expression especially at the BCP transcriptional (BCP double mutations; A1762T/G1764A) and translational sites (Kozak sequences mutants, nt1809-1812), and PC initiation (1814-1816). PC translational stop codon (G1896A with C1858T) and post translational mutant gene (G1862T) were equally examined [14,15,16]. Additionally, nucleotide changes in other BCP functional genome regions including the TA rich genome regions (nt1741-1755, nt1758-1762, nt1771-1775, and nt1788-1795) as well as initiation sites of PC mRNA at nt1788-1791 and pregenomic RNA at nt1817-1821 were also examined [15,16]. All mutant variants detected were compared among the two study groups.

#### DNA sequence accession numbers

The HBV BCP/PC sequences generated in this study have been deposited into National Centre for Biotechnology Information (NCBI) GenBank under accession number MT680878-MT680883.

### Statistical analysis

Accuracy and completeness of questionnaires were checked, data were double entered to reduce data entry errors and later merged. All categorical data and median (interquartile range, IQR) of continuous variables were compared by Chi-square and Mann-Whitney tests, respectively using SPSS version 25.0 (IBM, USA). The Optical density(O.D.) values of HBeAg and HbeAb were also compared between the study groups by Mann-Whitney test using GraphPad Prism (version 8.4.2, 2020). For all statistical analyses, P-value less than was considered significant at 95% CI.

## Results

### Demographic characteristics of study participants

The mean age of HIV/HBV co-infected patients was 40 years while HBV mono-infected patients was 30.5 years (p = 0.000). Overall, female had the highest seroprevalence of HBV infection among the two studied groups (62.5%, p = 0.003) (Table 1).

**Table 1.**
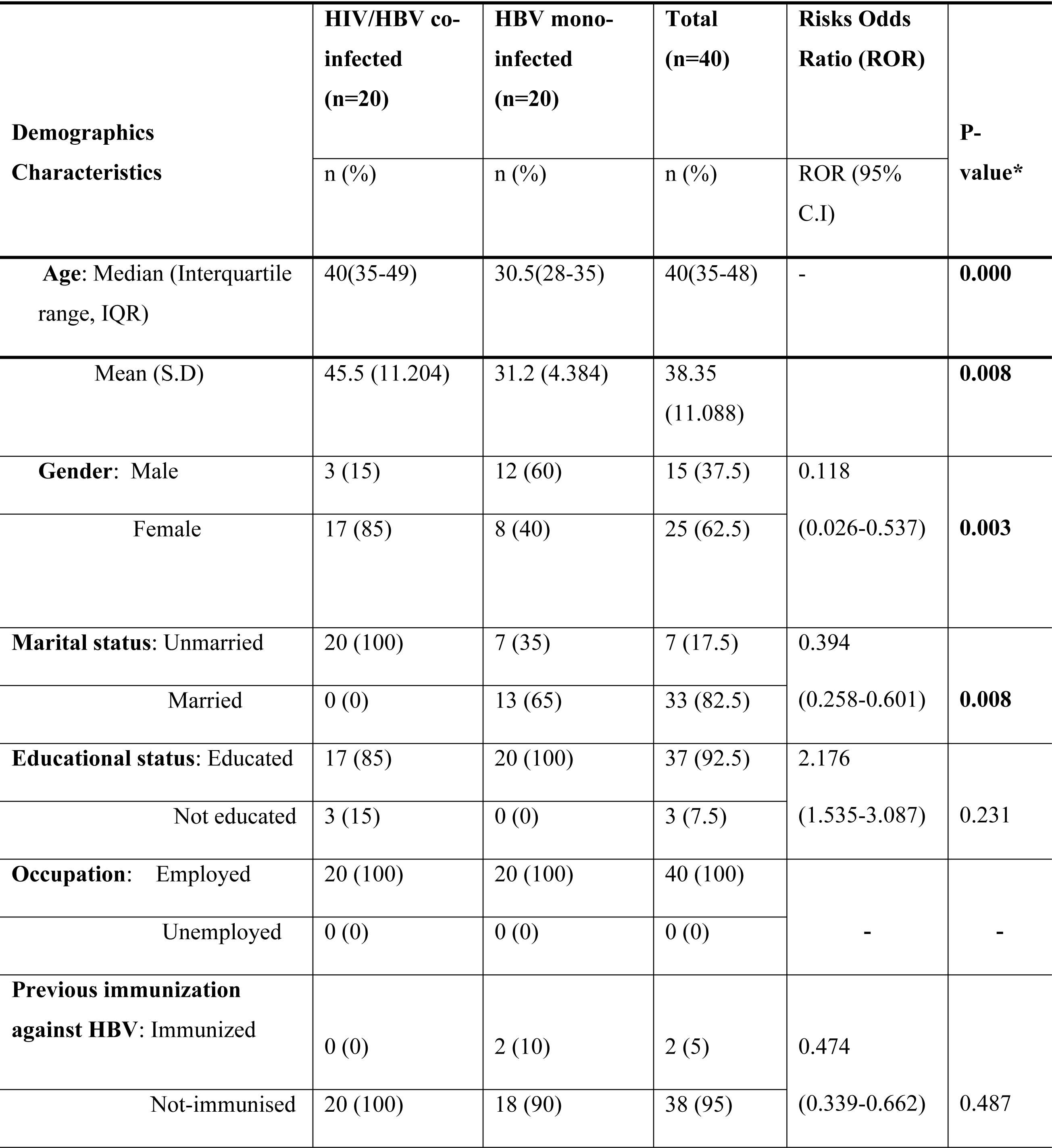
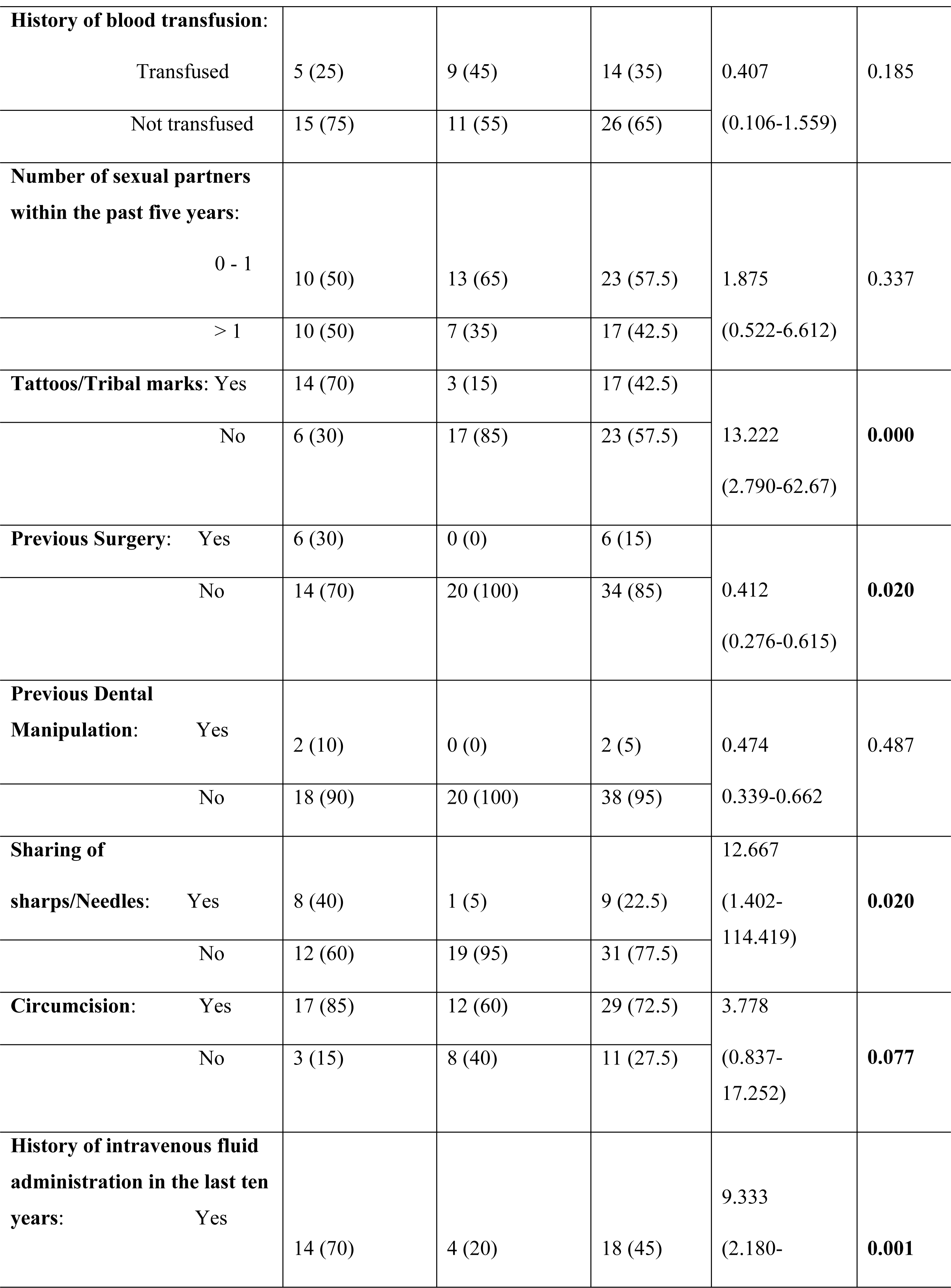
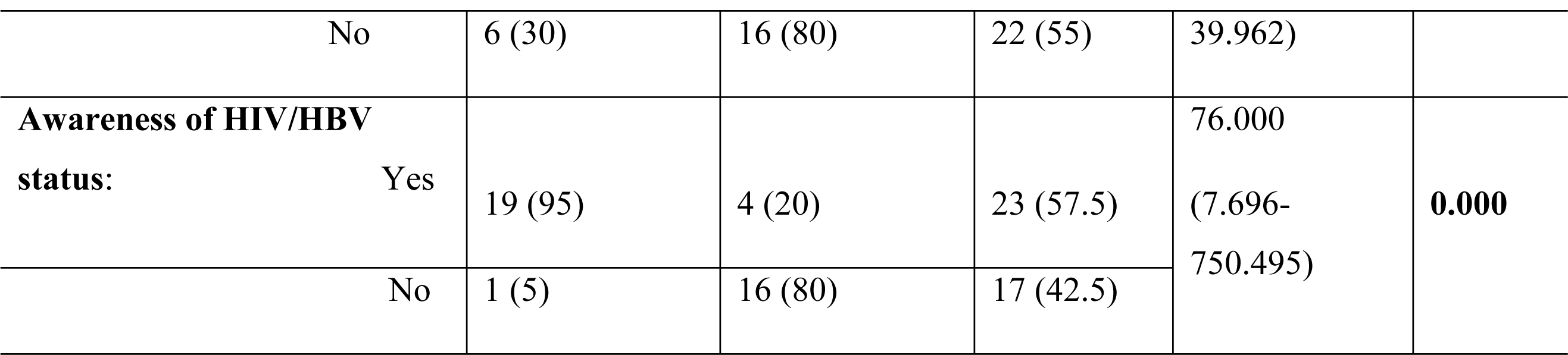
Demographic characteristics of study subjects.

### HBeAg and HBeAb ELISA screening and Genotypic analysis

Frequency of HBeAg and HBeAb varied across the two studied groups. Highest (23%) frequency of HBeAg was observed in the HIV/HBV co-infected group. Also, higher (60%) frequency of HBeAb was observed among the HBV mono-infected group (Fig 1). Genotyping analysis of the sequences (3 exploitable sequences) that were positive for S-gene specific nPCR showed that all the sequences belonged to genotype E (Fig 2).

**Figure 1:**
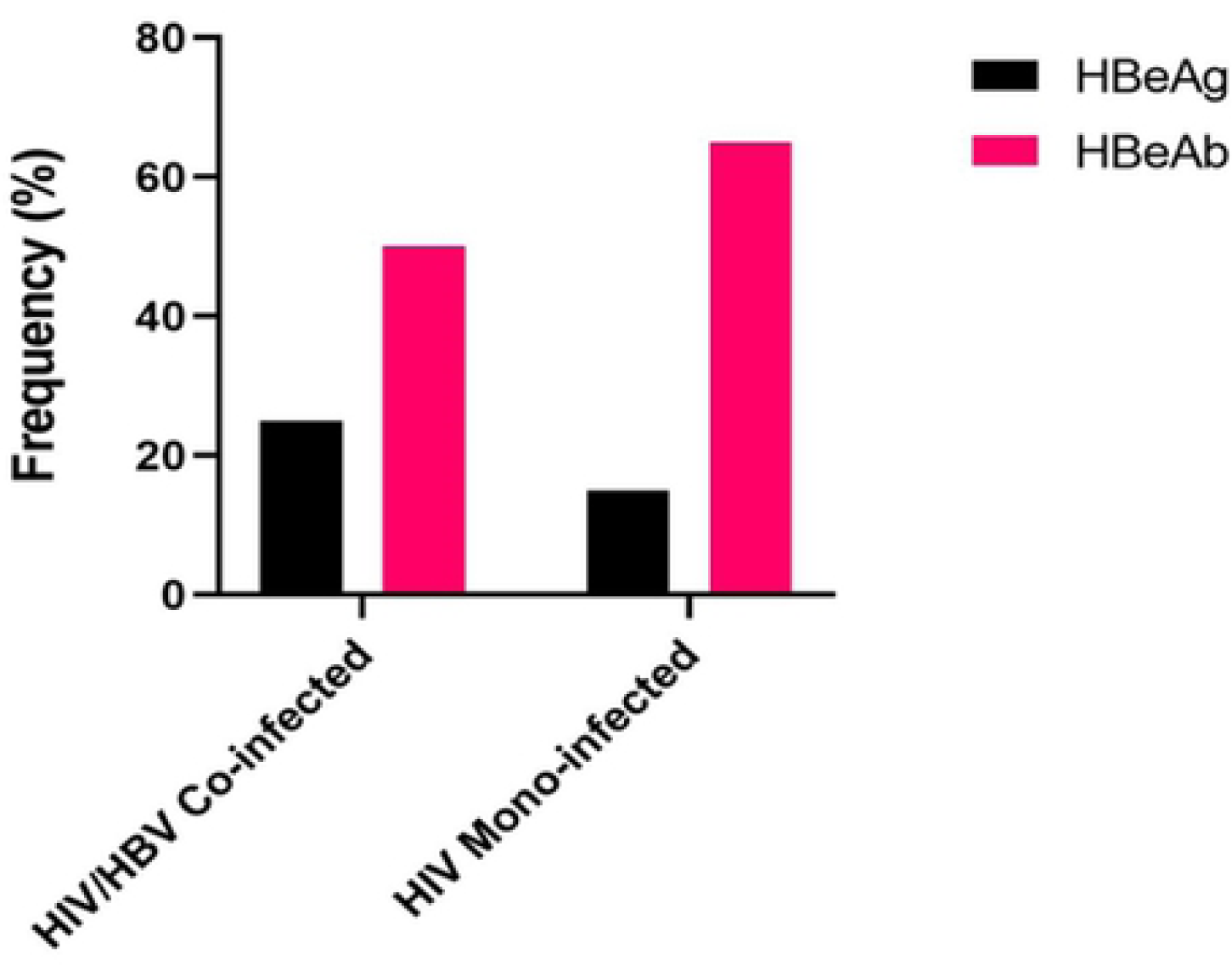
Frequency of HBeAg and HBeAb among HIV/HBV co-infected and HBV mono-infected patients in Ile-ife, Nigeria.

**Fig 2.**
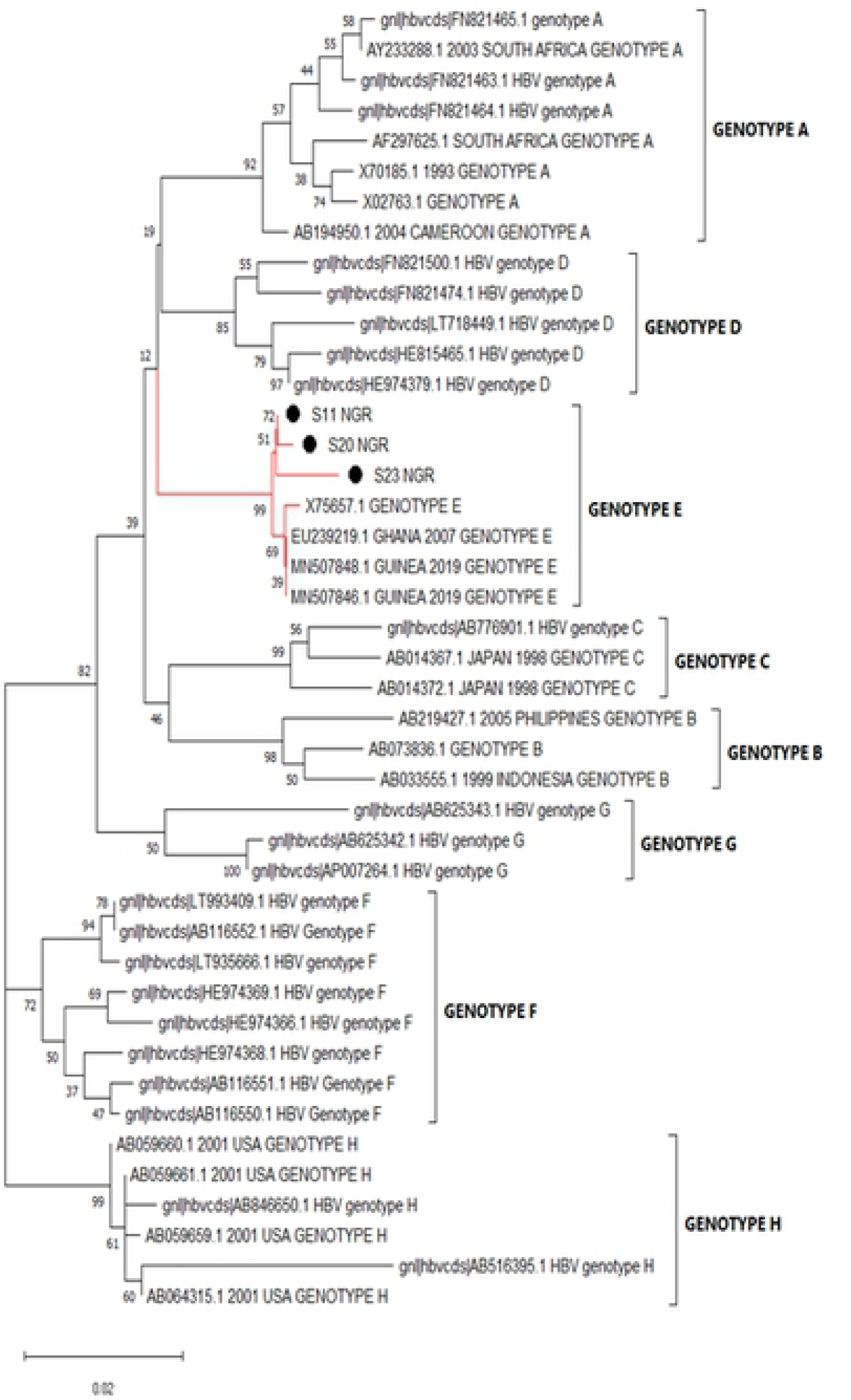
Phylogenetic analysis of HBV S gene sequences. The evolution ary history was inferred using Ille Negihbour-Joining method. The optimal tree with the sum of branch length= 0.39698535 is shown. The percentage of replicate trees in which the associated taxa clustered together in the bootstrap test (1000 replicates) are shown next to the branches. The tree is drawn to scale, with branch lengths in the same units as those of the evolutionary distances used to infer the phylogenetic tree. The evolutionary distances were computed using the Maximum Composite Likelihood method and are in the units of the number of base substitutions per site. This analysis involved 43 nucleotide sequences. All ambiguous positions were removed for each sequence pair (pairwise deletion option). There were a total of 400 positions in the final dataset. Evolutionary analyses were conducted in MEGAX.

### Variation analysis of the BCP/PC region

Of the 40 samples tested by BCP/PC region-specific nested PCR, 17 (42.5%) of the samples yielded the expected 306 bp. Specifically,12 and 5 samples from patients with HBV/HIV co-infection and HBV mono-infection respectively as shown by macrogen preliminary gel report in Fig 3. However, out of the 17 samples, seven (7) samples had low concentration/ volume and were not sequenced while the remaining ten samples were sequenced. Of the ten samples, sequence data of four samples were not exploitable due to the presence of multiple peaks. Sequence data of the remaining exploitable six (6) sequences were used for the BCP/PC mutational analysis.

**Fig 3:**
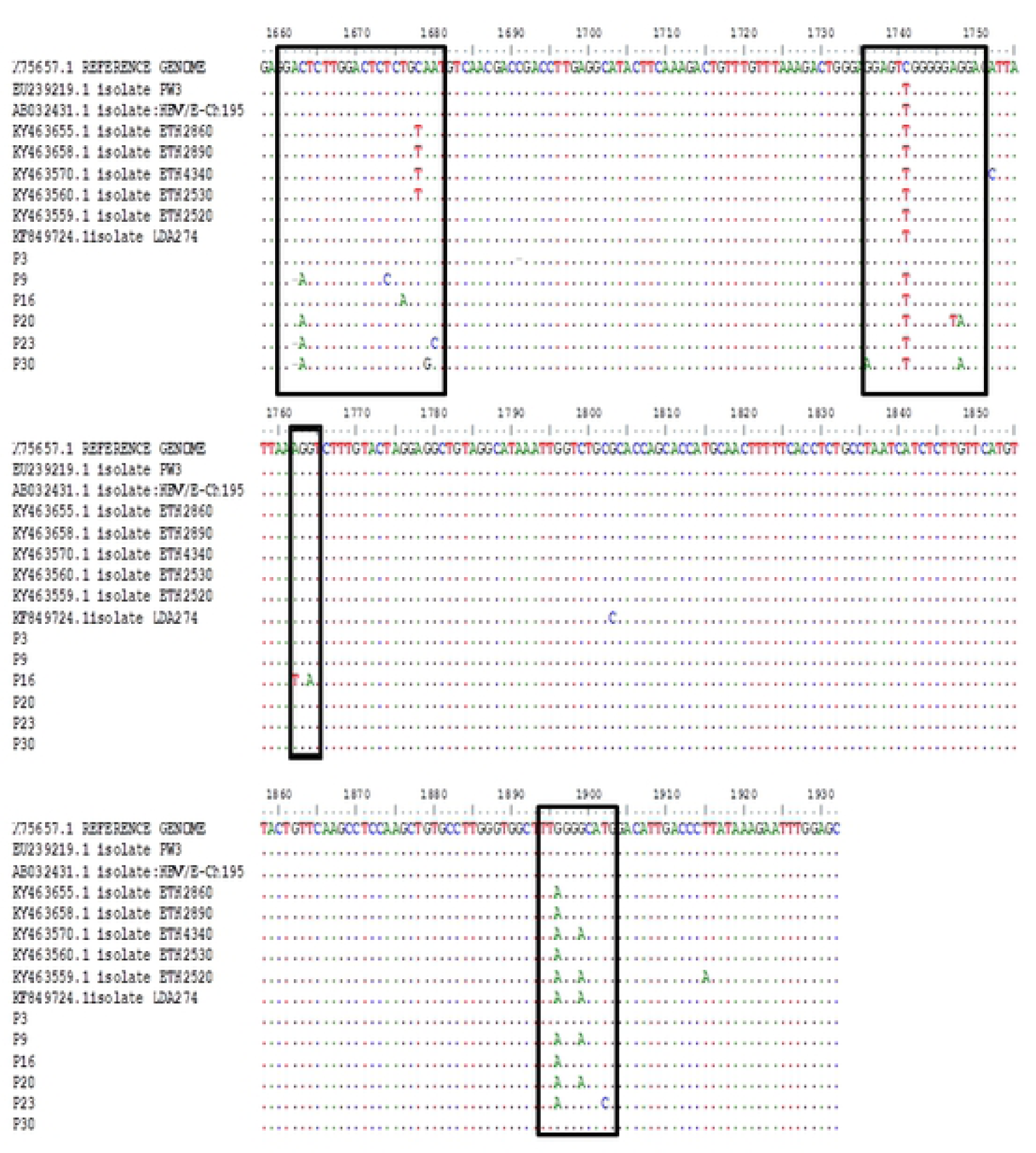
Preliminary gel electrophoresis report. Mutation patterns from patient samples against reference genomes from NCBI

Overall, 5(83.3%) of the 6 exploitable sequences after analysis showed various BCP/PC mutations (Figure 3). Interestingly, only 1 (16.6%) sequence from an HIV/HBV co-infected patient showed the BCP transcriptional (double mutation; A1762T/G1764A) mutant (Figure 3). There was no significant Kozak sequence (nt1809-1812) or PC initiation codon (nt1814-1816) mutants detected (Figure 3). Analysis of the PC translational stop codon showed 4(66.6%) having the G1896A mutants while 2 (33.3%) had G1899A mutants (Fig 4). None of the sequences in this study had the PC post-translational mutant variant (G1862T) (Fig 5).

**Figure 4:**
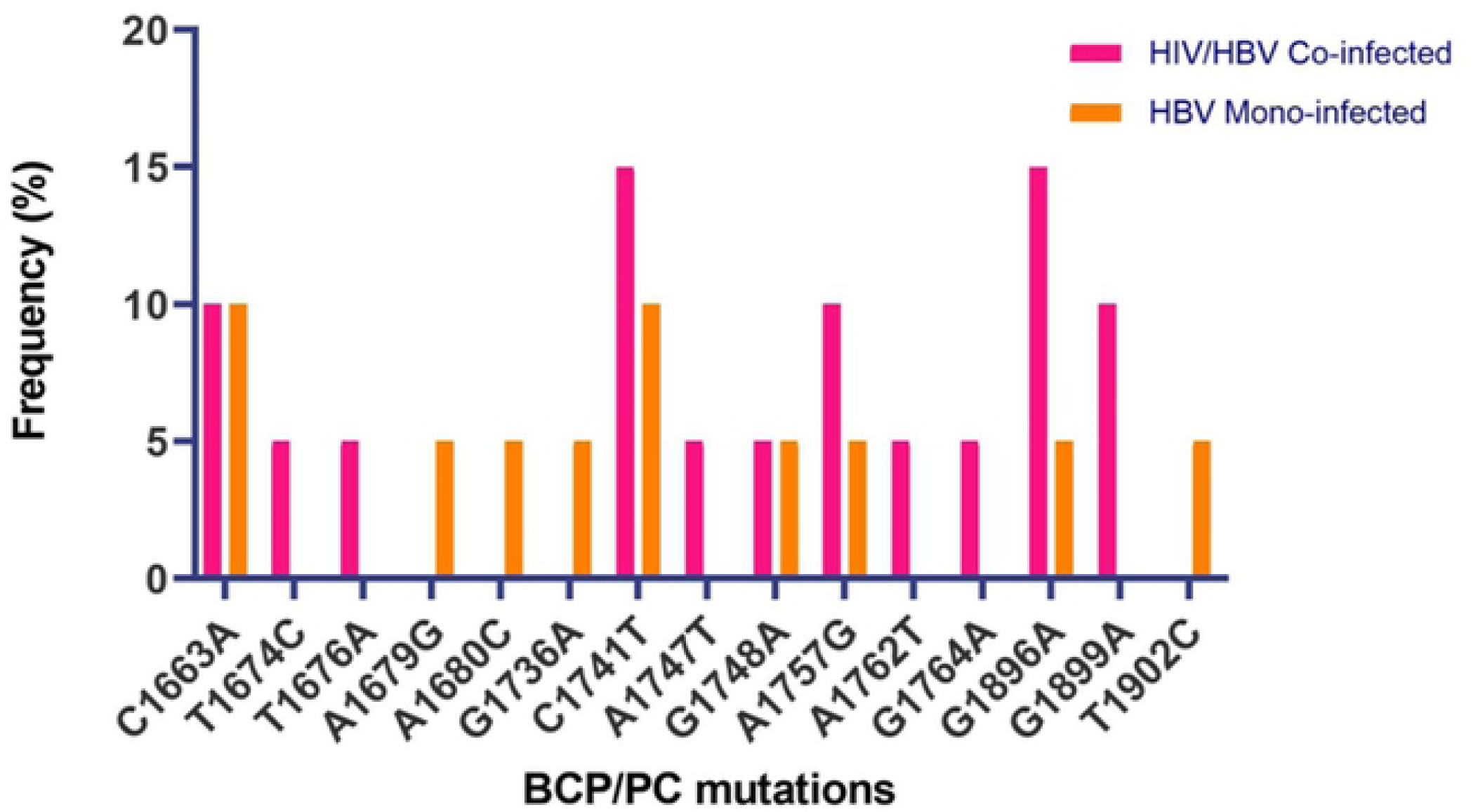
Mutation patterns from patient samples against reference genomes from National Centre for Biotechnology Information. prevalence pattern of HBV basal core promoter/precore region molecular mutant strains

**Figure 5:**
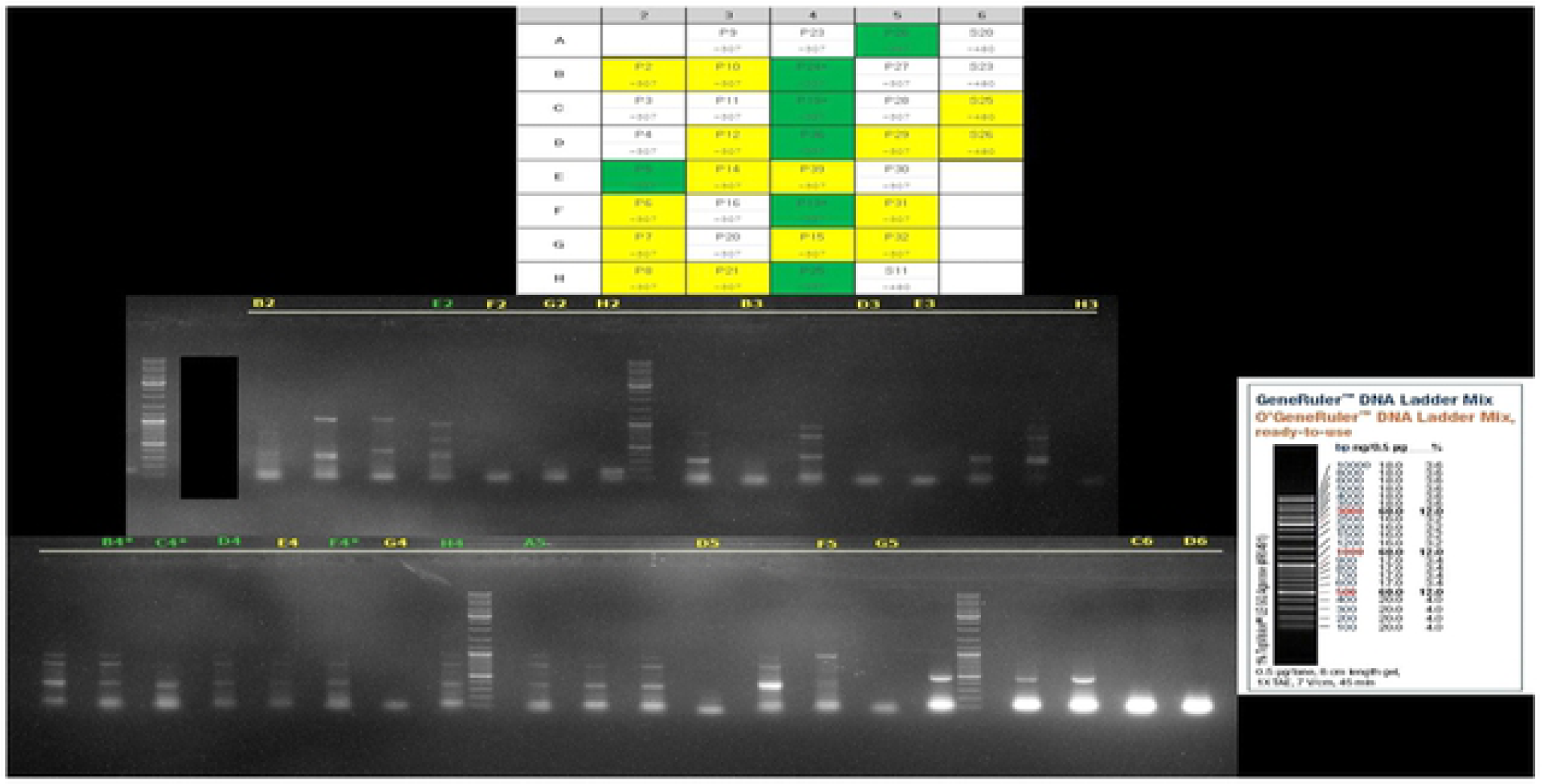
Prevalence pattern of HBV basal core promoter /precore region molecular mutant strains. Macrogen preliminary gel detntphoreas report Green colour sarnies -Too low concentration, <5ul Yellow colour samples - No band seen on gel electrophoresis Samples with an asterisk. Low concentration and low volume samples White coloured samples: possible to proceed for sequencing

## Discussion

The prevalence of HBsAg among people living with HIV/AIDS (PLWHA) using ELISA from our study was found to be 8%. This falls in the lower limit of the national range of HBV prevalence in HIV/HBV co-infected individuals in Nigeria. The prevalence of HBV in pools of HIV positive patients ranges from 10% to 70% in different subgroups of HIV positive Nigerians [17] and in fact, Nigeria has the highest variation in the prevalence of HBV among HIV/HBV co-infected individuals worldwide^17.^ This lies within the regional prevalence for HBV in HIV/HBV co-infected individuals in sub-Saharan Africa ranging from 0% to 28.4% in a review by Singh et al [18]

Serological findings from our study showed reduced frequency of HBeAg was observed in the HBV mono-infected group in comparison to the HBV/HIV co-infected cohort. This is not unexpected as documented evidence showed that HIV infected patients have reduced rate of spontaneous clearance of HBeAg [18]. HBV mono-infected individuals mount higher quantitative and qualitative HBeAb response in comparison to HIV/HBV co-infected individuals [19] with resultant effective clearance of HBeAg [19]. Also, this study confirms a higher prevalence of HBV pre-core region mutant strains in HIV/HBV co-infected cohort with resultant reduction in the secretion of HBeAg and invariably inadequate HBeAb response.

The emergence of HBeAg negative variants of HBV in HIV co-infected individuals have profound implication on the diagnosis, management and prognosis of this subset of individuals [20]. This includes reduced HBeAb production with associated increased rate of viral replication, persistence of HBV infection, development of chronic liver diseases and rapid progression to endstage liver diseases [20]. Currently, little is known about HBV pre-core region genomic heterogeneity in HIV/HBV co-infection in Nigeria and sub-Saharan Africa. In this study, we broadened the available evidence of pre-core/core region genomic variability among HIV/HBV co-infected patients in Nigeria and assessed the difference of mutation pattern with the HBV mono-infected patients in Nigeria. We found the HBV pre-core region gene variants C1663A, T1676A, A1679G, A1680C, G1736A, A1747T, G1748A, A1762T, G1764A, G1896A, G1899A, and T1902C. Interestingly, the HIV/HBV co-infected population had the highest frequency of core/pre-core mutant strains for the C1741T and G1896A (Figure 4). G1896A mutation is the commonest precore region mutation globally [20,21] which this study confirmed, and further established the high frequency of these HBV pre-core region mutant strains in Nigeria.

It is worth mentioning that earlier studies had established a positive correlation between G1896A mutation and progressive liver diseases [20,21,22]. Further, G1896A mutation among others has been reported to enhance HBV DNA replication even after HBeAg seroconversion which results in prolonged hepatic inflammation [21,22] However, some other researchers were of the contrary opinion. However, Kim et al were of the opinion that other factors including HBV genotype, geographical location, race, ethnicity, host immune competence and co-infection like HIV, HCV might be a determining factor in the progression of liver diseases in patients with G1896A mutation [23]. From our own study, we suggest that the presence of HIV/HBV co-infection might worsen the progression of liver diseases due to synergistic effect of HIV on the evolution of HBV pre-core region mutant strains.

From this study, we were also able to identify basal core promoter mutant strains-A1762T and G1764A from a patient among the cohorts of HBV/HIV co-infected individuals. These HBV variants are associated with cytoplasmic localization of intracellular HBcAg with associated active necroinflammation of the hepatocytes [24]. These mutations act as early predictors for progression to hepatocellular carcinoma [24]. Hepatocellular carcinoma (HCC) is a unique malignancy with a very high case fatality rate of approximately one [25]. Identifying these mutations in this specific patient provides a very important window of opportunity to screen and monitor the patient to prevent development of HCC.

Only 3 of the 40 HBV DNA amplicons yielded exploitable S gene sequenced data. The phylogenetic analysis of the three sequenced data showed that all the three HBV variants were genotype E. This genotype circulates in West Africa [26]. HBV exhibits genotype-specific pathogenesis; for instance, genotype C and D have higher risk of liver cirrhosis and hepatocellular carcinoma while genotypes A and B have better response to interferons than genotypes C and D [26]. HBV genotype E shows good response to nucleoside analogues [26]

Limitations of the study included inadequate sample size yield as we were only able to identify 20(8%) HBsAg positive samples from the cohort of 250 HIV positive patients screened. This sample size appears inadequate to make a general inference to the study population. In addition, only a few of the samples sequenced (6) were exploitable for BCP/PC mutational analysis and therefore affected the ability to generalize the findings. From our study, being the first in Nigeria to identify hepatitis B virus pre-core region molecular mutant strains among cohorts of HBV/HIV co-infected and HBV mono-infected individuals, future research should focus on the need to look at the role of HAART in the evolution of the HBV pre-core region molecular mutant strains especially among subsets of drug-naive and drug experienced HIV/HBV co-infected individuals in Nigeria. Furthermore, there is the need to follow up the lone patient from the study with C1762T and G1764A for proper management and prevention of development of hepatocellular carcinoma.

## Conclusion

This study has broadened the available evidence of hepatitis B virus precore region molecular mutants among cohorts of HIV/HBV co-infected and HBV mono-infected individuals in Nigeria and Africa.. We therefore recommend that HIV/HBV co-infected patients be routinely screened for hepatitis B virus precore region molecular strains to improve their patient outcome.

## Acknowledgements

We wish to express our profound gratitude to Professor Olaniyi Onayemi, Professor Dennis Ndububa, Dr Patricia Gorak-Stolinska, Dr Folarin Onikepe and Dr Tope Faleye for their expert consultation, advice, encouragement and contribution towards the success of the research work.

